# The breadth of protection mediated by anti-N2 neuraminidase antibody responses relies on cross-reactive and not cross-inhibiting antibodies

**DOI:** 10.64898/2026.02.02.703223

**Authors:** Laura Amelinck, Anouk Smet, Tine Ysenbaert, Koen Sedeyn, William Warren, Raul Gomila, Thorsten U. Vogel, Xavier Saelens, João Paulo Portela Catani

## Abstract

Neuraminidase (NA) inhibition (NAI) titers have been identified as an independent correlation of protection against influenza. Few studies, however, have investigated the breadth of NA-based immune protection. Previously, we have reported that N2 NAs derived from human H3N2 viruses that circulated between 2009 and 2017 can be subdivided into four antigenic groups. Here, we immunized mice with recombinant soluble tetrameric NA from H3N2 strains representing those four antigenic groups or passively transferred N2 NA immune serum into naïve mice to evaluate the breadth of protection against a heterologous HxN2 influenza virus challenge. We show that the breadth of protection goes beyond the breadth of NAI but still requires the presence of cross-reactive antibodies. Interestingly, in the absence of cross-reactive antibodies, the immunization of DBA/2J mice with heterologous NA was associated with an early onset of disease upon challenge with the reassortant HxN_ind11_ or H2N2 A/Singapore/1/1957.

## Introduction

Hemagglutinin (HA) and neuraminidase (NA) are the major surface antigens of influenza virions. These two glycoproteins exert a complementary role in the influenza virus replication cycle: while HA binds to sialic acids, an essential step for virion entry, NA cleaves off terminal sialic acid residues [1]. This balanced activity of HA and NA enables influenza virions to penetrate the mucus barrier, rich in decoy receptors, lining the respiratory tract thereby enabling access to sialic acid receptors on the apical side of respiratory epithelial cells. Additionally, NA activity is crucial for the release of newly formed viral progeny from the infected cell [2,3]. NA inhibiting antibodies are thought to prevent or attenuate influenza by impairing these steps in the viral life cycle.

Traditionally, the protective humoral immune response against influenza virus infection has been attributed to the ability of antibodies to neutralize the virus by blocking attachment to sialic acid receptors present on the surface of susceptible cells. The presence of such neutralizing antibodies is typically evaluated with a hemagglutination inhibition assay performed with sera from immunized or convalescent subjects, red blood cells and influenza virions. Hemagglutination inhibition (HAI) is accomplished by anti-HA antibodies and constitutes the gold standard correlate of protection against flu. Despite HA being the major target of influenza vaccines, studies from the 1970s already pointed out the protective effect of NA as an immunogen [4,5]. More recent studies have confirmed those early observations and established that anti-NA antibodies are an independent correlate of protection against influenza [6–8]. The protective effect of NA inhibition has also been demonstrated in a post-exposure prophylaxis: oseltamivir treatment protected close contacts of influenza virus-infected individuals [9]. Despite NA’s protective potential, its immunogenicity is suboptimal in conventional split inactivated influenza vaccines and NA is absent in recombinant hemagglutinin vaccines [10,11].

Monoclonal antibodies can inhibit NA activity by occluding the catalytic site. Such antibodies prevent NA-mediated cleavage of small substrates, such as 2′-(4-Methylumbelliferyl)-α-D-N-acetylneuraminic acid (MUNANA, a fluorogenic NA substrate) or NA-STAR (1,2-dioxetane derivative, a chemiluminescent substrate). Antibodies that bind NA distal to the catalytic site can also interfere with NA activity, particularly when the substrate is a complex sialic acid-containing glycoprotein such as fetuin; these antibodies impede access of the NA catalytic site to fetuin by steric hinderance [12]. Finally, NA-specific antibodies that do not inhibit NA activity can still protect mice from challenge with influenza virus. This protection relies on Fc effector functions, which are also involved in protection mediated by antibodies with weak NAI activity [13–15].

Previously, we determined the antigenic diversity of NA from human H3N2 viruses circulating from 2009 to 2017 by assessing ferret and mouse immune sera against a panel of HxN2 reassortant viruses in a fetuin-based NAI assay [14]. The resulting NAI patterns revealed four antigenic groups (AGs) among the selected human H3N2 NAs [16]. Here, using representative strains from each AG, we evaluated in mice the breadth of protection conferred by active immunization with recombinant NA (tetNA) or passive transfer of NA immune sera against challenge influenza virus. Cross-protection was found to correlate with the presence of cross-reactive antibodies determined by ELISA.

## Results

### Pathogenicity of HxN2 strains in mice

Six NA representatives spanning the four AGs were selected for use as NA immunogens or as the NA component of a HxN2 reassortant challenge viruses (Figure 1a) [16]. To circumvent the poor pathogenicity of H3N2 strains in mice, we established challenge models using H6N2 reassortant viruses, in which H6 is derived from A/mallard/Sweden/81/2002 (H6N1) (hereafter, Hx). In an initial attempt to establish an HxN2 challenge model in mice, female BALB/c mice were inoculated with 10-fold serial dilutions of HxN_A/Perth/16/2009nib-64 (Per09)_. None of the mice, however, exhibited sign of disease or weight loss, even at the highest dose (8.2×10^3^ PFU per mouse) (Supplementary Figure 1a). Since DBA/2J mice are reported to have increased susceptibility to influenza virus infection, the experiment was repeated in this strain using a 3-fold serial dilutions of HxN_A/Singapore/infimh-16-0019/2016 (Sin16)_, HxN_Per09_, HxN_A/Texas/50/2012 (Tex12)_, HxN_A/Kansas/14/2017 (Kan17)_, HxN_A/Helsinki/823/2013 (Hel823)_, and HxN_A/Indiana/08/2011 (Ind11)_ (Figure 1b-g) [17]. All HxN2 reassortant viruses were pathogenic in DBA/2J mice, except HxN_Sing16_ (Figure 1 and Supplementary Figure 1b).

**Figure 1.**
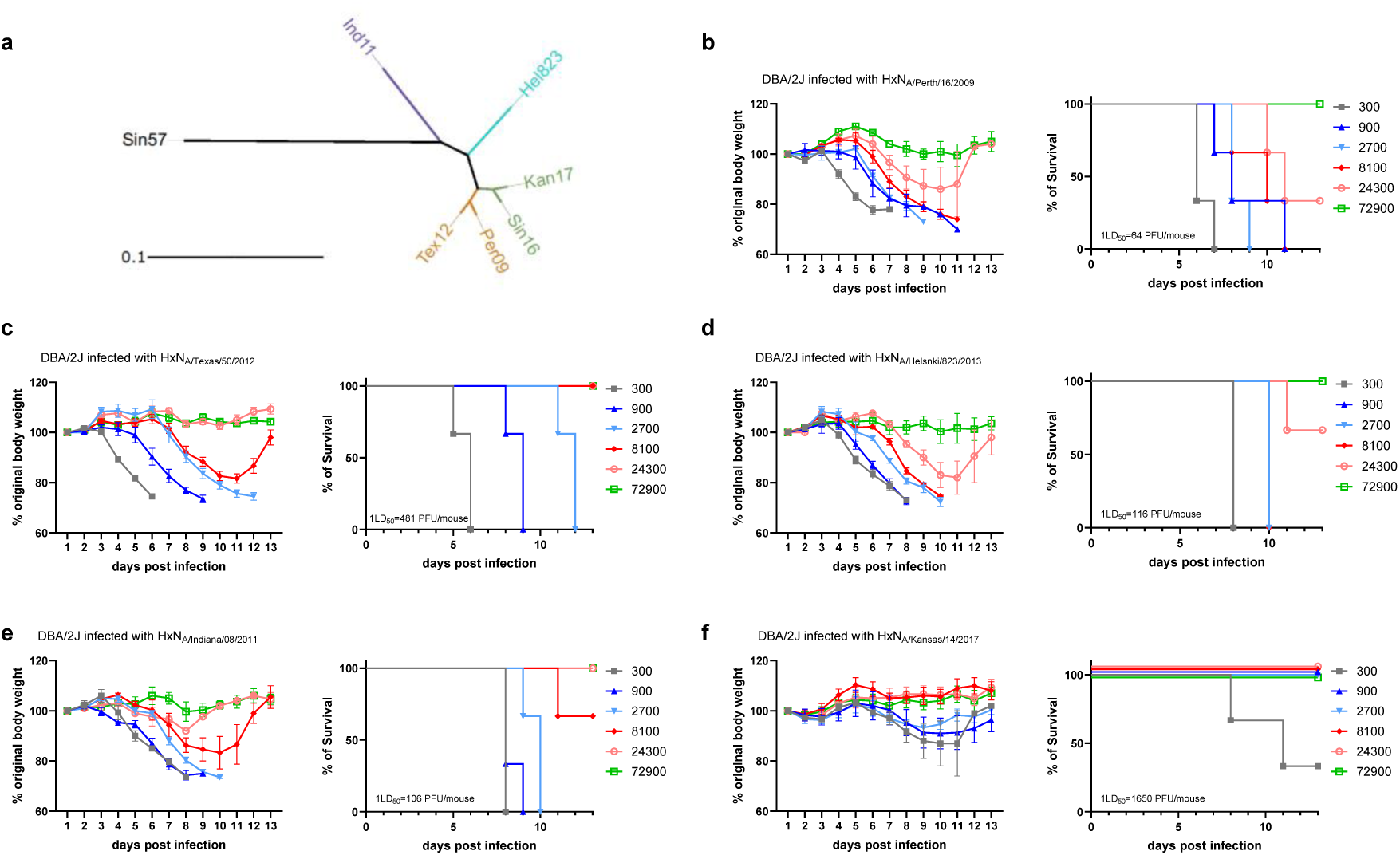
Antigenicity and pathogenicity of HxN2 reassortant viruses in DBA/2J mice. (**a**) Phylogenetic tree of NA amino acid sequence depicting color-coded AGs. Tex12 and Per09 belong to AG1 (dark yellow); Sin16 and Kan17 to AG2 (green); Hel823 to AG3 (Blue), and Ind11 to AG4 (purple). (**b-f**) The relative body weight, survival and LD50 of DBA/2J mice inoculated with (**b**) HxNA/Perth/16/2009nib-64, (**c**) HxNA/Texas/50/2012, (**d**) HxNA/Helsinki/823/2013, (**e**) HxNA/Indiana/08/2011, and (**f**) HxNA/Kansas/14/2017. The LD50 values were calculated using the Reed–Muench method.

### Protection induced by immunization with NA from distinct antigenic groups against HxN_A/Perth/16/2009nib-64_

Given that A/Perth/16/2009 (H3N2) has been used in a controlled human infection model, we first determined whether immunization with NAs from distinct AGs would protect mice against HxN_Per09_ [18].

Mice were primed and boosted with tetrabrachion-stabilized recombinant NA (tetNA), in a two-week interval; serum was collected, and viral challenge was performed 2 weeks after the boost. ELISA endpoint titers using microtiter plates coated with NA from A/Perth/16/Perth09 (Per09 NA) showed that homologous immunization induced the highest serum IgG titers (Figure 2a). NA from A/Texas/50/2012 (Tex12 NA), which belongs to the same AG as Per09, also induced high titers of cross-reactive antibodies (Figure 2a). Mice immunized with NA from A/Singapore/infim-16-0019/2016 (Sin16 NA) had high titers of cross-reactive antibodies, although these were significantly lower than those induced by homologous immunization (Figure 2a). Finally, mice immunized with NAs from A/Helsinki/823/2013 (Hel823 NA) and A/Indiana/08/2011 (Ind11 NA) had significantly lower cross-reactive antibodies titers against Per09 NA (Figure 2a).

**Figure 2.**
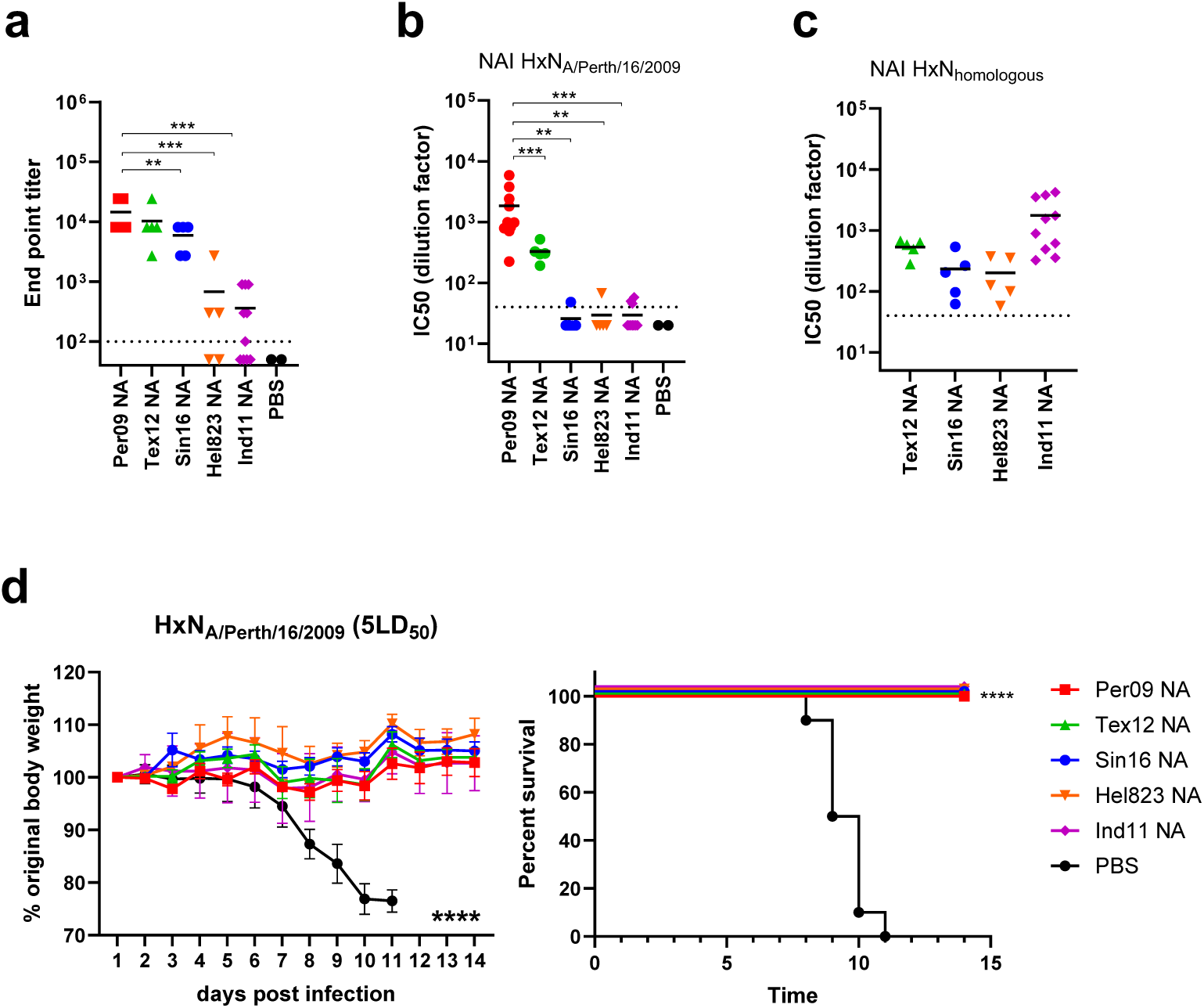
Serology and cross-protection of mice immunized with heterologous tetNAs and challenged with HxNPert09. DBA/2J mice were prime-boosted in a two-week interval regimen with 0.1 µg of AF03-adjuvanted Perth09, Tex12, Sing16, Hel823, or Ind11 tetNA. (**a**) ELISA was performed with sera obtained two weeks after the boost using Perth09 tetNA that was captured in nickel-coated plates. (**b**) NAI was determined using HxN2 HxNA/Perth/16/2009 and HxN2s with homologous NA used in the immunization (**c**). Two weeks after the boost mice were challenged with 5LD50 of the reassortant HxNPer09. The graphs show the percentage of initial body weight and mortality (**d**). Statistical comparison of the ELISA endpoint dilution titers was performed using Mann-Whitney test. Differences between NAI titers were analyzed using Welch’s t test. Data points in a-c are from individual mice except for the PBS group, which represent pools of 5 mice each, and horizontal solid lines represent means. Dashed lines represent the limit of detection. Data points in d represent averages of individual mice and error bars SEM. Statistical analysis of relative body weight changes was assessed by an approximate F-test and survival using Mantel-Cox rank. Data in d and e are pooled from 2 independent experiments except for NA Tx12, NA Sin16, and NA Hel823, which are from 1 experiment. Dashed lines represent the limit of detection. \*\**P*<0.01,\*\*\**P*<0.001, \*\*\*\**P*<0.0001 compared with the PBS immunized group.

NAI titers of mouse immune sera were also measured against HxN_Per09_. As expected, the highest NAI titers were observed in mice immunized with Per09 NA. NAI titers were also detected in the group immunized with the heterologous Tex12 NA, but these were significantly lower than those in the homologous group. Mice immunized with N2 from different AGs—Sin16 NA, Hel13 NA, and Ind11 NA—had low or undetectable NAI titers against HxN_Per09_ (Figure 2b). To exclude the possibility that the reduced ELISA titers and NAI against Per09 reflected failed immunization, we also performed NAI assays using HxN2s homologous to each antigen. The homologous NAI titers confirmed that all groups responded as expected (Figure 2c).

Two weeks after boost, mice were challenged with 5LD_50_ of HxN_Per09_. All mice immunized with tetNA were protected and only the PBS control group exhibited weight loss and mortality (Figure 2d).

### Protection induced by immunization with NA from A/Perth/16/2009 against HxN2 viruses with NA derived from distinct antigenic groups

Given that immunization with distinct NAs conferred protection against challenge with HxN_Per09_, we next evaluated the reciprocal question: whether immunization with Per09 NA could protect against challenge with HxN2s viruses bearing NAs from distinct AG. Four HxN2 viruses (HxN_Tex12_, HxN_Kan17_, HxN_Hel823_, and HxN_Ind11_) were used to challenge mice that had received homologous NA (matching the challenge virus NA; not available for HxN2_Kan17_), heterologous Per09 NA, or PBS.

The reciprocal experiment confirmed the antigenic similarity between Per09 and Tex12: immunization with Per09 NA induced ELISA titers against Tex12 NA that were similar to homologous Tex12 immunization (Figure 3a). NAI titers of sera from Per09 NA-immunized mice against HxN2 Tex12 were similar to those from Tex12 NA-immunized mice against HxNPer09, confirming bidirectional cross-reactivity between these antigens (Figures 3b and 3c). Consistent with these serologic data, both Per09- and Tex12 NA-immunized mice were protected against challenge with 5LD_50_ of HxN_Tex12_ (Figure 3d).

**Figure 3.**
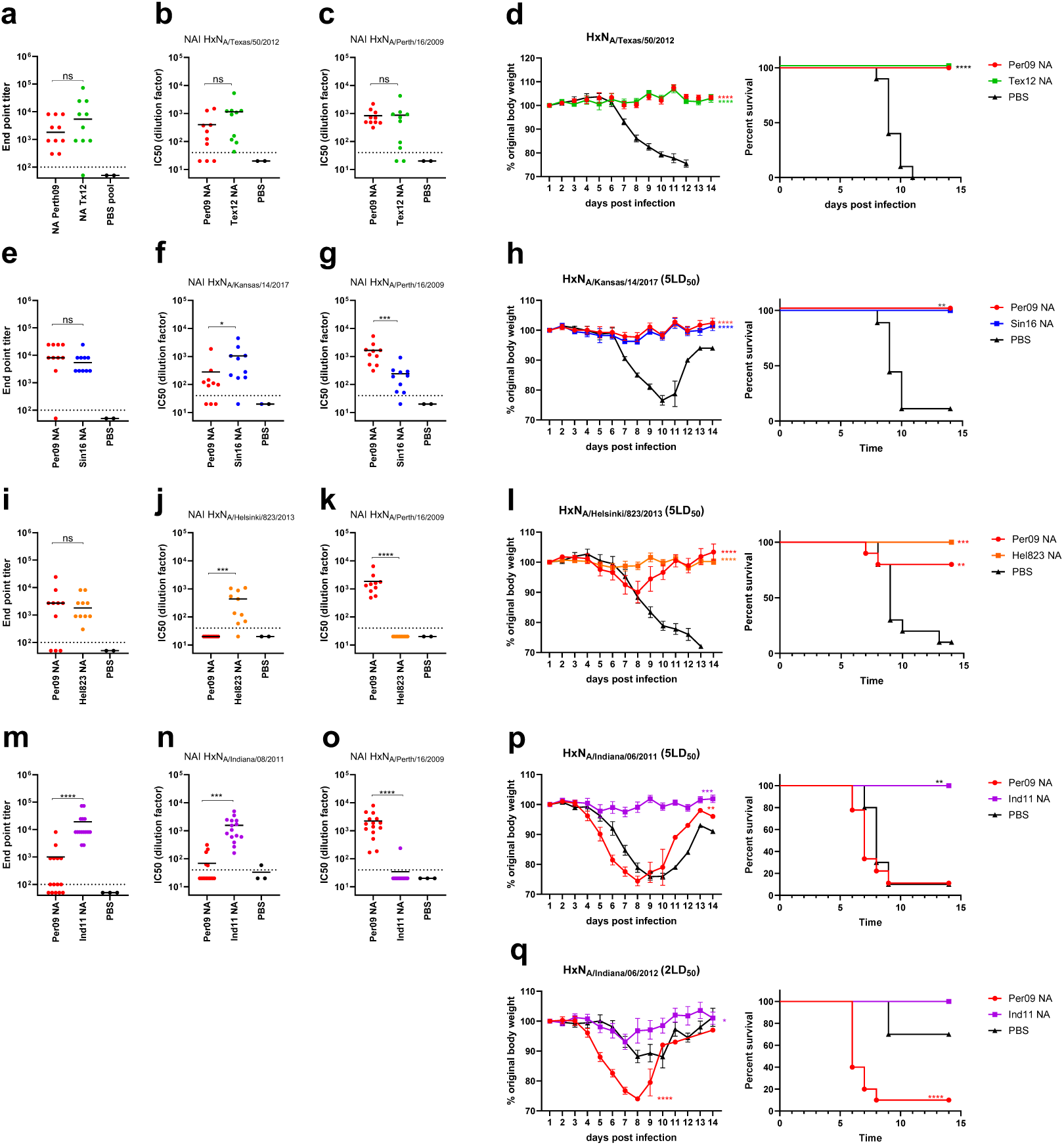
Serology and cross-protection of mice immunized with Per09 tetNA and challenged with heterologous HxNx reassortants. DBA/2J mice (n = 5 per group per experiment) were prime-boosted in a two week interval with 0.1 µg of AF03-adjuvanted tetNA or PBS as indicated. ELISA and NAI assays were performed with sera isolated two weeks after the boost immunization. (**a**) ELISA was performed using Tex12 tetNA that was captured in nickel coated plates. (**b**) NAI of immune sera determined against HxNTex12 and (**c**) HxNPer09. (**d**) Two weeks after the boost, mice were challenged with 5 LD50 of HxNTex12. The graphs show the percentage of initial body weight and mortality. (**e**) ELISA was performed against Sin16 tetNA. NAI was determined against (**f**) HxNKan17 (**g**) and HxNPer09. (**h**) Relative body weight and mortality of mice challenged with 5LD50 of the HxNKan17 virus. (**i**) ELISA was performed against Hel823 tetNA. NAI determined against (**j**) HxNHel823 and (**k**) HxNPer09. (**l**) Relative body weight and mortality of mice challenged with 5LD50 of HxNHel823 virus. (**m**) ELISA against Ind11 tetNA. NAI determined against (**n**) HxNInd11 or (**o**) HxNPer09. Relative body weight and mortality of mice challenged with (**p**) 5 or (**q**) 2 LD50 of HxNInd11. Statistical comparison of the ELISA endpoint dilution titers was performed using Mann-Whitney test. Differences between NAI titers were analyzed using Welch’s t test. Statistical analysis of relative body weight changes was assessed by an approximate *F*-test and statistical analysis of survival using Mantel-Cox rank. Dashed lines represent the limit of detection. All experiments were repeated once and the data pooled for analysis. \**P*<0.05, \*\**P*<0.01,\*\*\**P*<0.001, \*\*\*\**P*<0.0001 compared with the PBS immunized group, or as indicated.

We next selected Sin16 NA because it belongs to a distinct AG to Per09 and was a WHO-recommended vaccine strain in 2018-19 northern hemisphere (NH) season. Since HxN_Sin16_ is not pathogenic in mice, HxN_Kan17_ was used for challenge studies, as it belongs to the same AG as Sin16 NA. This AG is characterized by the substitutions S245N/S247T, and P468H in NA[16,19]. Anti-Sin16 NA IgG titers measured two weeks after the boost were similar in mice immunized with Per09 NA and Sin16 NA (Figure 3e). NAI titers against the HxN_Kan17_ challenge strain were higher in mice immunized with Sin16 NA; however, detectable NAI titers were observed in sera from Per09 NA immunized mice (Figure 3f). Conversely, NAI titers against HxNA_Per09_ were higher in Per09 NA-immunized mice, but detectable titers were also observed in Sin16 NA-immunized mice (Figure 3g). Immunization with Per09 and Sing16 NA equally protected mice against challenge with 5LD_50_ of HxN_Kan17_ (Figure 3h).

We also investigated protection induced by Per09 NA against HxN_Hel823_. Although isolated in 2013, this antigenically distinct N2 is genetically similar to neuraminidases from H3N2 viruses that circulated more than a decade earlier. Immunization with Per09 NA induced cross-reactive IgG titers against Hel823 NA comparable in magnitude to those obtained with homologous Hel823 NA immunization (Figure 3i). Despite these cross-reactive antibodies, sera from Per09 NA-immunized mice did not inhibit the NA activity of HxNA_Hel823_ (Figure 3j). Similarly, sera from mice immunized with Hel823 NA did not show detectable NAI against HxN_Per09_ (Figure 3k). Despite the lack of cross-reactive NAI titers, evidence of heterologous cross-protection after challenge was observed. Mice immunized with heterologous Per09 NA showed transient weight loss, they were similarly protected compared with mice immunized with homologous Hel823 NA (Figure 3i). This finding suggests that cross-reactive, non-inhibiting NA-specific antibodies can contribute to protection against heterologous NA challenge.

We next evaluated whether immunization with Per09 NA would confer protection against HxN_Ind11_. Ind11 NA belongs to a distinct AG and is derived from a swine origin H3N2 virus isolated from humans. Immunization with Per09 NA induced low or undetectable cross-reactive ELISA titers against Ind11 NA, whereas all mice immunized with the homologous Ind11 NA showed detectable titers (Figure 3m). Similarly, NAI titers against HxN_Ind11_ were mostly absent or at lower levels in Per09 NA immune sera compared with Ind11 NA-immunized mice (Figure 3n). In line with this observation, NAI antibodies against HxN_Per09_ were detected in Per09 NA-immunized mice but not in the Ind11 NA immunized mice (Figure 3o). Only mice that had been immunized with Ind11 NA were protected against mortality and morbidity when challenged with 5 LD_50_ of HxN_Ind11_; an early onset of morbidity was observed in the group immunized with heterologous Per09 NA (Figure 3p). To better evaluate this earlier onset of morbidity, we performed a similar experiment with a lower challenge dose (2LD_50_), which confirmed the observation (Figure 3q).

### Protection mediated by NA immune response is transferable by sera, even in the absence of NA inhibiting antibodies

Cross-protection against influenza can be mediated by cross-reactive antibodies or cellular responses directed against conserved antigens [2,20–22]. To evaluate whether protection observed after NA immunization relied on antibody responses, we performed serum transfer experiments in which mice received anti-NA immune sera representing the 4 NA AGs (Per09, Sin16, Hel823, and Ind11). Mice were then challenged with 2LD_50_ of HxN2 viruses carrying NA from the respective AGs.

First, serum was generated by immunizing DBA/2J mice with 1 µg of AF03-ajuvanted NA in a prime-boost regimen with a two-week interval. Two weeks after the boost, immune serum was prepared, pooled by group, and analyzed by ELISA and NAI to confirm seroconversion prior to transfer to naïve mice. Mice that had received serum against Per09, Sin16, and Hel823 were protected against challenge with HxN_Per09_, whereas Ind11 NA immune serum recipient were not (Figure 4a). All transferred sera had detectable ELISA endpoint titers against Per09 NA (Figure 4b), whereas NAI titers against the HxN_Per09_ challenge virus were detectable only in homologous Per09 NA and heterologous Sin16 NA immune sera (Figure 4c). When challenged with 2LD_50_ of HxN_Kan17_, sera from the 4 AGs were protective against weight loss, while sera from naïve mice were not (Figure 4d). Survival curves did not differ between the groups challenged with 2LD_50_ of HxN_Kan17_(Figure 4d). All NA immune sera contained cross-reactive antibodies against Sin16 NA, however, only Ind11 NA immune serum had undetectable NAI titers against HxN_Sin16_ (Figures 4e and 4f). Kan17 and Sin16 NA share 98.5% amino acid sequence identity and belong to the same NA AG.

**Figure 4.**
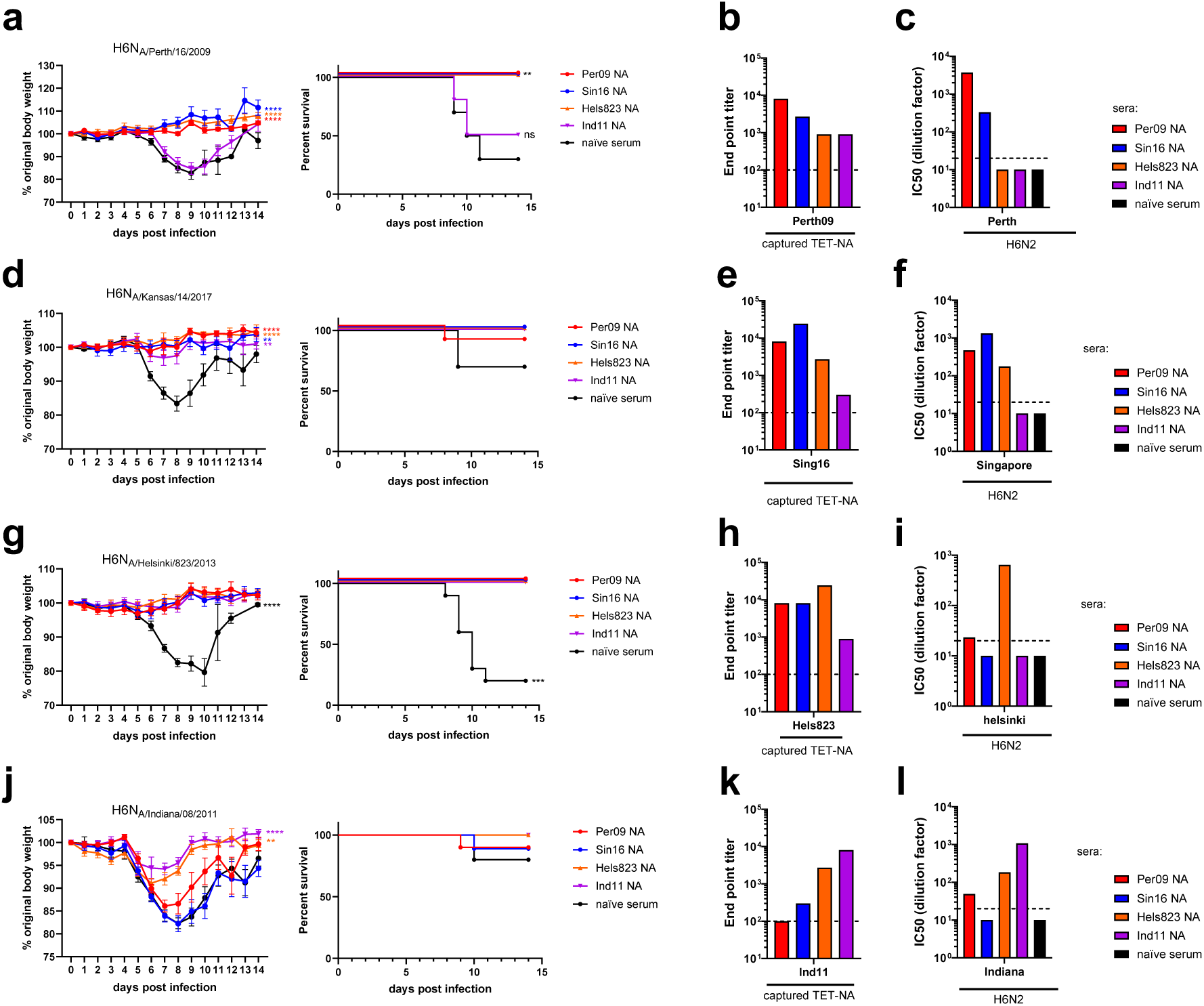
Mice that received immune serum with NA cross-reactive IgG are protected against heterologous HxNx challenge. Serum used for passive transfer was obtained from DBA/2J mice that had been immunized in a prime-boost regimen with 1 µg of AF03-Adjuvanted tetNA or PBS (naïve serum). One day after serum transfer, mice were challenged with 4 different HxN2 viruses. (**a**) Relative body weight and survival of mice challenged with 2 LD50 of HxNPer09. (**b**) ELISA performed using Per09 tetNA that was captured in nickel coated plates. (**c**) NAI determined against the HxNPer09 challenge virus. (**d**) Relative body weight and survival of mice challenged with 2 LD50 of HxNInd11. (**e**) ELISA performed using Sin16 tetNA that was captured in nickel coated plates. (**f**) NAI determined using HxNSin16 virus, a reassortant with NA that belongs to the same AG as the NA of A/Kansas/14/2017. (**g**) Relative body weight and survival of mice challenged with 2 LD50 of HxNHel823. (**h**) ELISA performed using Hel823 tetNA that was captured in nickel coated plates. (**i**) NAI determined against HxNKan17. (**j**) Relative body weight and survival of mice challenged with 2 LD50 of HxNInd11. (**k**) ELISA performed using Ind11 tetNA that was captured in nickel coated plates. (k) NAI determined against HxNInd11. The graphs in panels a, d, g, and j show the percentage of initial body weight and mortality. Statistical analysis of relative body weight changes was analyzed as repeated measurements using method of residual maximum likelihood (REML) and survival using Mantel-Cox rank. Challenge experiments were performed twice and data from the 2 independent experiments were pooled for analysis. Dashed lines represent the limit of detection. \*\**P*<0.01; \*\*\*\**P*<0.0001 compared with the PBS immunized group.

Mice that received any of the anti-NA immune sera were protected against challenge with HxN_Hel823_ (Figure 4g). Although cross-reactive antibodies against Hel823 NA were detected in all sera, only homologous sera had detectable NAI titers against HxN_Hel823_ (Figures 4h and 4i). Finally, most mice survived the challenge with 2LD_50_ of HxN_Ind11_, and only mice that had received homologous Ind11 NA or Hel823 NA immune serum had reduced weight loss compared with naïve serum recipients (Figure 4j). High titers of Ind11 NA reactive antibodies were detected in Ind11 NA and Hel823 NA immune sera, whereas titers in Per09 NA and Sin16 NA immune sera were near the limit of detection (Figure 4k). Ind11 NA and Hel823 NA immune sera had high NAI titers against HxN_Ind11_, whereas Per09 NA and Sin16 NA immune sera had low or undetectable NAI titers. It is important to consider that the tetrabrachion zipper, although not immunodominant, is immunogenic, and immunization with recombinant tetNAs induces detectable IgG titers against this zipper domain (Supplementary figure 2)[23].

### Immunization with Per09 NA does not protect mice against A/Singapore/1/57 H2N2 challenge

We also assessed whether immunization with Per09 NA would protect against challenge with H2N2 A/Singapore/1/1957 (Sin57). Sin57 was not pathogenic in BALB/c mice and was pathogenic to DBA/2J mice only at a very high inoculum (LD_50_ of 1.8×10^5^ PFU/mouse, Supplementary figure 3). Therefore, we performed mouse adaptation of Sin57. After 10 passages in mouse lungs and MDCK cells, the mouse-adapted virus exhibited increased pathogenicity, with 1LD_50_ of 4.6×10^2^ PFU/mouse. The mouse-adapted Sin57 carried a G103S and K422R substitution in HA and a G111D substitution in NA.

Mice were immunized with homologous Sin57 NA or heterologous Per09 NA. Two weeks after the boost, the mice were challenged with 5LD_50_ of mouse-adapted Sin57. Neither homologous nor heterologous NA immunization prevented infection, as all mice experienced at least transient weight loss following challenge. However, all mice immunized with homologous Sin57 NA survived, whereas mice immunized with heterologous Per09 NA were not protected (Figure 5a). We also observed an early onset morbidity and mortality trend in challenged mice immunized with Per09 NA. To further evaluate this observation, immunized mice were challenged with 0.5LD_50_ of mouse-adapted Sin57.

**Figure 5.**
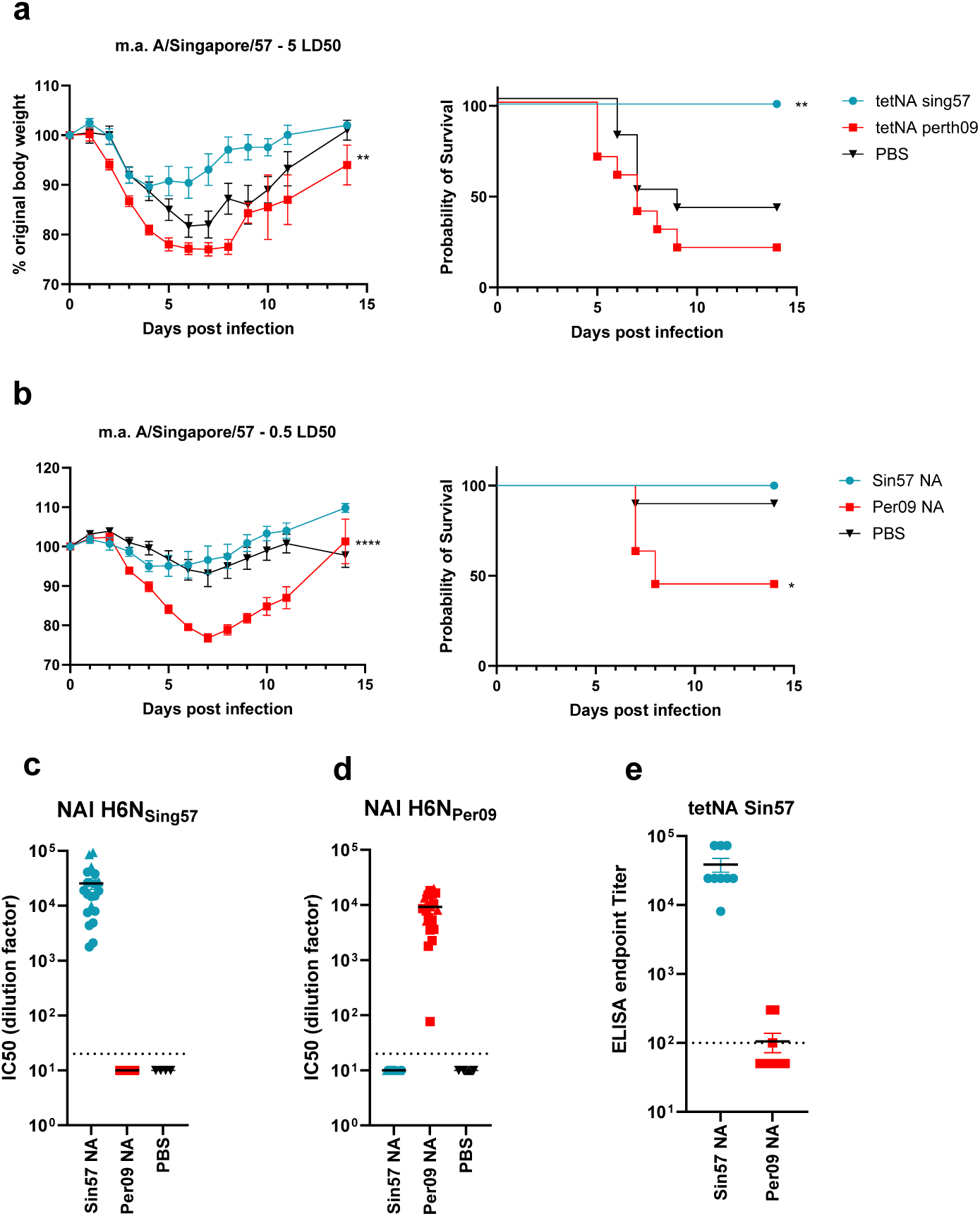
Immunization with Per09 NA does not protect mice against A/Singapore/1/57 H2N2 challenge. Mice were immunized in a prime-boost regimen with 0.1 µg of AF03-adjuvanted tetNA or PBS control. (**a**) Relative body weight and survival of mice challenged with 5LD50 of mouse-adapted Sin57. (**b**) Relative body weight and survival of mice challenged with a dose of 0.5LD50 of mouse-adapted Sin57. NAI determined against (**c**) HxNSin57 or (**d**) HxNPer09. (**e**) IgG titers determined by ELISA against Sin57 NA. Statistical analysis of relative body weight differences was analyzed as repeated measurements using method of residual maximum likelihood (REML) and survival using Mantel-Cox rank. Challenge experiments were performed twice and data from the 2 independent experiments were pooled for analysis. Dashed lines represent the limit of detection. \**P*<0.05; \*\**P*<0.01; \*\*\*\**P*<0.0001 compared with the PBS immunized group.

While mice immunized with homologous Sin57 NA or PBS control experienced a minor transient weight loss, the group immunized with heterologous Per09 NA displayed increased morbidity and mortality (Figure 5b). No cross-NAI antibodies were detected against HxN_Sin57_ and HxN_Per09_ (Figure 5c and d, respectively). In addition, most Per09 NA-immunized mice did not have detectable Sin57 NA cross-reactive ELISA titers.

## Discussion

Although NAI titers are an independent correlate of protection, NA has not yet been implemented as a standardized influenza vaccine component [24]. This is due in part to the labile nature of NA and to our limited understanding of NA antigenicity relative to HA. Previously, we defined 4 distinct NA AGs in human H3N2 viruses isolated between 2009 and 2017 [16]. Here, by active immunization of mice with tetrabrachion-stabilized N2 NAs representative of these 4 groups, we demonstrate that protection against an HxN2 virus challenge can occur in the absence of NA inhibiting antibodies.

To circumvent the poor pathogenicity of recent human H3N2 viruses in mice, we used HxN2 reassortants bearing H6 HA from A/mallard/Sweden/81/2002, N2 NAs from selected human H3N2 viruses, and the internal segments of the PR8 laboratory strain [25–27]. Whereas inoculation of BALB/c mice with these HxN2 viruses did not induce signs of disease, DBA/2J mice proved susceptible to most HxN2 challenges, with LD_50_ values ranging from 64 to 1650 PFU/mouse. Still, the HxN_Sin16_ was not pathogenic, and HxN_Kan17_ was pathogenic at doses 3.4 to 25 times higher than the other tested HxN2 viruses. Sin16 NA and Kan17 NA belong to AG 2, whose NAs exhibit a higher Km, potentially resulting in a suboptimal functional balance between H6 and N2 [16,28].

After establishing challenge models with HxN2 representatives of the 4 NA AGs, we performed a series of active immunizations using AF03-adjuvanted NAs, followed by challenge. Our first question was whether antigenically distant N2 NAs would provide protection against HxN_Per09_. Interestingly, despite not inducing detectable cross NAI titers, immunization with Sin16 NA, Hel823 NA, and Ind11 NA fully protected mice against challenge with heterologous HxN_Per09_. These immunizations induced cross-reactive antibodies against Per09 NA. This result was partially recapitulated by serum transfer experiments, in which all anti-NA sera except Ind11 NA protected against virus challenge. In line with this, active immunization with heterologous Per09 NA also protected against heterologous HxN_Tex12_, HxN_Kan17_, and HxN_Hel823_ challenge even in the absence of cross-NAI antibodies. Thus, the presence of cross-reactive antibodies sufficed to protect mice against heterologous HxN2 virus challenge, a finding recapitulated by serum transfer.

Intriguingly, mice immunized with Per09 NA and challenged with heterologous HxN_Ind11_ showed an early onset morbidity and mortality. This observation was not recapitulated by serum transfer experiments, in which only homologous Ind11 NA and heterologous Hel823 NA immune sera with detectable NAI were able to confer protection.

To assess whether the breadth of N2 NA-based immune protection could be expanded beyond the previously defined AGs, mice immunized with Per09 NA or homologous NA were also challenged with H2N2 Sin57. Similar to the HxN_Ind11_ setting, immunization with heterologous Per09 NA induced an early onset of morbidity and mortality after challenge with mouse-adapted Sin57, which was most evident at a sublethal dose. Per09 NA immunization induced neither cross-inhibiting nor cross-reactive antibodies against H2N2 NA, consistent with increased divergency of Per09NA antigen and N2 from A/Singapore/1/1957 challenge (Supplementary figure 4) and with the consequent lack of observed protection.

The early onset of morbidity and mortality of the Per09 NA-immunized mice challenged with HxN_ind11_ was not observed in the serum transfer experiments, suggesting a cellular mechanism may be at play. This is supported by the H2N2 challenge results, where similar early onset of morbidity occurred even in the absence of cross-inhibiting and cross-reactive antibodies. Further research is required to elucidate the mechanisms underlying early disease in NA-immunized mice challenged with heterologous N2 NA-containing influenza viruses. Our data does not support this as general phenomenon, rather it is restricted to HxN_ind11_ and H2N2 A/Singapore/01/1957 strains.

It is worth noting that some of the detected cross-reactive antibodies may be directed against the tetrabrachion domain. This zipper is immunogenic, yet not immunodominant, and may be less accessible in the capture ELISA used to determine NA-specific IgG titers (Supplementary figure 2). Overall, our data suggests that heterologous protection mediated by N2 NA-specific antibodies does not rely on cross-inhibiting antibodies but rather on cross-reactive antibodies.

While a number of clinical studies have identified NAI as an independent correlate of protection, with reported correlations to reduced viral shedding and shorter duration of symptoms [6,7,29–35], few studies have investigated whether this also applies to NA binding antibodies. Notably, a recent study reported that NA binding antibodies also correlate with protection from medically attended influenza caused by influenza A and B viruses [36]. To our knowledge, no study has yet compared the predictive value of NAI and NA-binding titers. Based on our mouse data, we hypothesize that protection predicted from NA-binding titers may be broader than that predicted from NAI titers. These findings further encourage the use of NA as a standardized vaccine antigen to improve protection against influenza.

## Materials and Methods

### Design and production of recombinant proteins

The coding information for tetNA fusion proteins (Per09, Tex12, Sin16, Hel823, Ind11, and Sin57 NAs) were cloned under the transcriptional control of the CMV promoter in the pCDNA3.4 plasmid with a CD5 secretion signal, an amino-terminal His-tag (except Sin57 which bears a strep-tag instead), and the thrombin cleavage signal followed by tetrabrachion and NA head domain (truncated stalk at position 75). Per09, Tex12, Sin16, Hel823, and Ind11 NAs were produced in CHO cells while Sin57 was produced in HEK293T cells. Recombinant proteins were purified by affinity followed by size exclusion chromatography as previously described [37].

### Viruses

The HxN2 reassortants were previously described in Catani *et al*., 2024 [16]. Briefly, these reassortant viruses express the targeted NA antigen, internal genes from A/Puerto Rico/8/1934 H1N1 (PR8), and the HA from A/mallard/Sweden/81/2002 (H6N1). All segments were cloned into a bidirectional transcription plasmid derived from pUC57 (Genscript) including RNA polymerase (Pol) I and Pol II promoters, as previously described [38]. The entire set of eight plasmids was used to transfect 293FT cells (Thermo Fisher Scientific) using Lipofectamine 2000 CD (Thermo Fisher Scientific). Twenty-four hours after transfection, Madin-Darby Canine Kidney Cells (MDCK-) ATL cells (ATCC) were added to the transfected cells in the presence of TPCK-treated trypsin (Sigma) to allow propagation of the rescued H6Nx viruses. Cell culture supernatants containing influenza virus were harvested 7 days post MDCK addition and blindly passaged in 8- to 10-day-old embryonated chicken eggs (Charles River Laboratories, Inc). Inoculated eggs were incubated at 37°C for 48 hr, then cooled to 4°C for 12 hr, prior to allantoic fluid harvest and clarification by low-speed centrifugation (3000 rpm, 20 min). High-yield stocks were generated by an additional passage in eggs as described above. Virus titers were determined by plaque assay on MDCK cells.

### ELISA

Anti-tetNA IgG titers were determined by capture ELISA using tetNAs in Pierce nickel-coated plates (cat. # 15442). Recombinant NA proteins were diluted to 0.5 µg/ml in DPBS (Life Technologies cat. # 14040-182). Then, 50 µl of the coating antigen solutions was added to each well and the plate was incubated at room temperature on a shaking platform (1h for capture ELISA and overnight for conventional ELISA). The wells of the plates were then washed three times with PBS-T (Sigma cat. # P3563-10PAK) and blocked for 1 hr with 1% BSA in DPBS. After blocking, wells were washed once with PBS-T and incubated with a threefold serial dilution, starting from a 1/100 dilution, of serum in DPBS with 0.5% BSA and 0.05% Tween-20 for 2 hr at room temperature on a shaking platform. Plates were then washed five times with PBS-T and incubated with a 1:5000 dilution of anti-mouse IgG-HRP (GE Healthcare cat. # NA931-1ml) in PBS with 0.5% BSA and 0.05% Tween20. The 3,3’,5,5’-tetramethylbenzidine (TMB) substrate (BD cat. # 555214) was added after three washes with PBS-T and the reaction was stopped after 5 min by addition of 50 µl of 1 M H_2_SO_4_. The optical density (OD) in each well was determined at 450 nm and, as a reference, 655 nm using an iMark Microplate Absorbance Reader (Bio-Rad). The end point titer was determined for each serum sample by scoring the dilution that resulted in an OD that was equal to or two times higher than the background OD obtained from pre-immune control sera dilution series.

### ELLA to determine NAI titers

Fetuin (Sigma cat. # F3385) was diluted into coating buffer (KPL cat. # 50-84-01) to a concentration of 25 µg/ml and 50 µl was added to the wells of Nunc MaxiSorp plates (Thermo Fisher cat. # 44-2404-21), which were subsequently incubated overnight at 4°C. The coated plates were then washed three times with PBS-T (Sigma cat. # P3563-10PAK) and incubated overnight with 25 μl of a dilution of HxNx that corresponds to 70% of the maximum activity of NA from the respective viruses as determined in the ELLA, and 25 µl of two-fold serial dilution of serum, starting at a 1/20 dilution, in sample buffer (1× MES VWR cat. # AAJ61979-AP: 20 mM CaCl_2_, 1% BSA, 0.5% Tween-20). Fetuin-coated plates were then washed three times with PBS-T and incubated for 1 hr with a solution of PNA-HRP (cat. # L6135-1MG, Sigma) at 5 μg/ml in conjugate diluent (MES pH 6.5, 20 mM CaCl_2_, 1% BSA). The plates were washed three times with PBS-T, TMB substrate was added, and then the plates were incubated for 5 min before the reaction was stopped by the addition of 50 µl of 1 M H_2_SO_4_. The optical density was measured at 450 nm and, as a reference, 655 nm in an iMark Microplate Absorbance Reader (Bio-Rad). Half maximum inhibitory concentrations (IC_50_) values were determined by non-linear regression analysis using GraphPad Prism software.

### Animal experiments

The mouse immunization experiments were conducted according to the Belgian legislation (Belgian Law 14/08/1986 and Belgium Royal Decree 06/04/2010) and European legislation on protection of animals used for scientific purposes (EU directives 2010/63/EU and 86/609/EEC). Experimental protocols were approved by the Ethics Committee of the Vlaams Instituut voor Biotechnologie (VIB), Ghent University, Faculty of Science (EC2021-045, EC2021-086 and EC2024-093). Female DBA/2J mice, aged 7-8 weeks, were purchased from Janvier (France). The mice were housed in a specified pathogen-free animal house with food and water ad libitum. Animals were immunized intramuscularly in the right quadriceps. Immunization was performed with 50 µl containing the recombinant protein. Protein based immunizations were adjuvanted with a 1:1 volume of AF03 (25 μl of antigen in PBS + 25 μl of AF03 per dose)[39].

Blood samples were obtained by tail vein puncture two weeks after boost immunization. For virus inoculation, the mice were anesthetized with 5% isoflurane and 50 μl of virus dilution was applied into both nostrils ensuring complete aspiration. Body weight was determined daily for 14 days after challenge. Mice were euthanized when they had lost more than 25% of body weight relative to the day of challenge.

For the serum transfer experiments, 100 μl of pooled sera prepared from mice that had been immunized with 1 μg of AF03-adjuvanted recombinant NA or PBS (with adjuvant), was injected intraperitoneally to DBA/2J mice one day prior challenge.

H2N2 Sin57 was mouse-adapted by 10 passages through lung and MDCK cells. For each passage, groups of 3 DBA/2J mice were inoculated with a 1/100 dilution of the viral stock and sacrificed 72 hr later to recover virus from the lungs. Lung homogenates were pooled and prepared in PBS using 7 ml tubes and a Precellys Evolution mechanical homogenizer, at 10000 RPM for two cycles of 2 min. The homogenate was clarified by centrifugation at 1,000 × g for 10 minutes at 4°C, then diluted 1:1000 prior to inoculation of MDCK cells. Infected cells were maintained in the presence of 2 µg/mL TPCK-treated trypsin (Sigma-Aldrich, T1426) until observation of cytopathic effect. The cell culture medium was then recovered, cleared by centrifugation and used at 1/100 dilution to inoculate naive mice. HxN2 challenge experiments were performed in BSL2 and experiments with A/Singapore/1/1957 in a BSL3 laboratory.

## Statistical analysis

Relative body weight values were analysed as repeated measurements using residual maximum likelihood (REML). Statistical analysis of survival was done using the log-rank (Mantel-Cox) test. ELISA titers were compared using the Mann-Whitney or Kruskal-Wallis test. NAI titers were compared using Welch’s t test or Kruskal-Wallis test. All statistical tests were performed using GraphPad Prism version 8.3.0. Statistical significance was considered when p < 0.05.

## Acknowledgements

A/Singapore/1/1957 virus was kindly provided by Jørgen de Jonge. We are grateful to the animal caretakers of the animal house at the VIB-UGent Center for Inflammation Research. The authors thank Jean-Sébastien Bolduc (Sanofi) for his critical review and proofreading of the manuscript.

## Author contributions

JPPC, TY, AS, KS and LA, performed experimental work. JPPC, TUV, and XS designed the experiments. JPPC and XS wrote the manuscript and are accountable for accuracy and integrity of the work. All authors proofread and critically reviewed the manuscript.

## Supplementary Figures

**Supplementary Figure 1.**
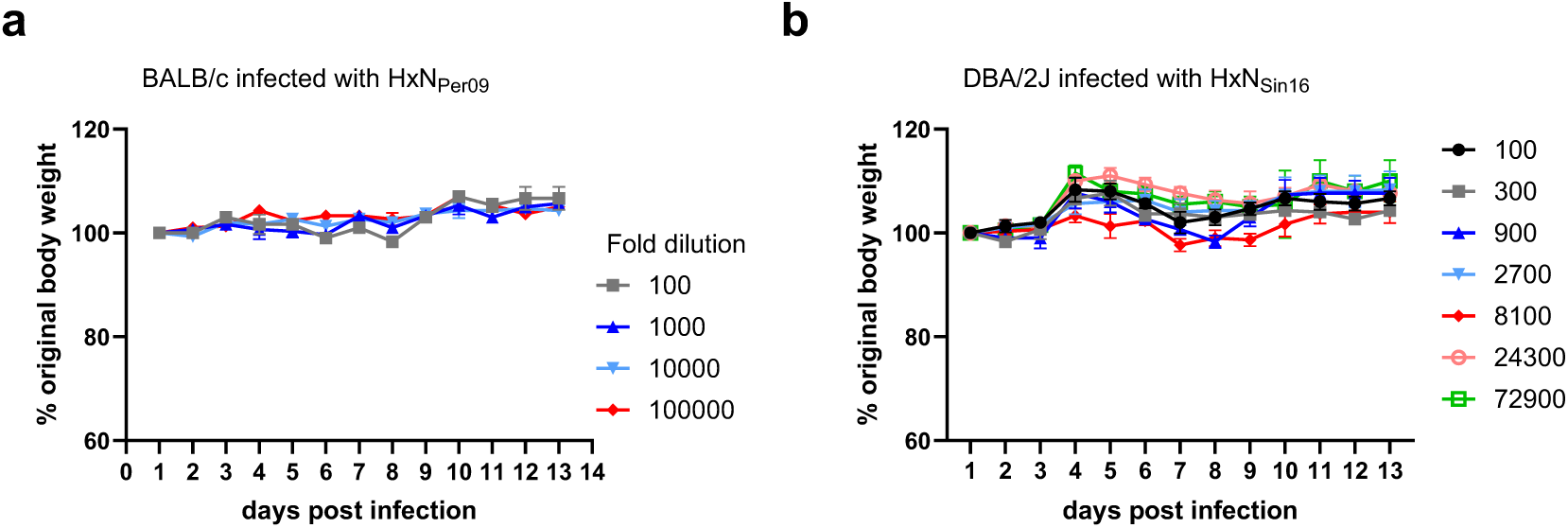
Pathogenicity of HxN2 reassortant viruses in mice. (**a**) Relative body weight of BALB/c mice inoculated with serial dilution of HxNPer09. (**b**) Relative body weight of DBA/2J mice inoculated with a serial dilution of HxNSin16.

**Supplementary Figure 2.**
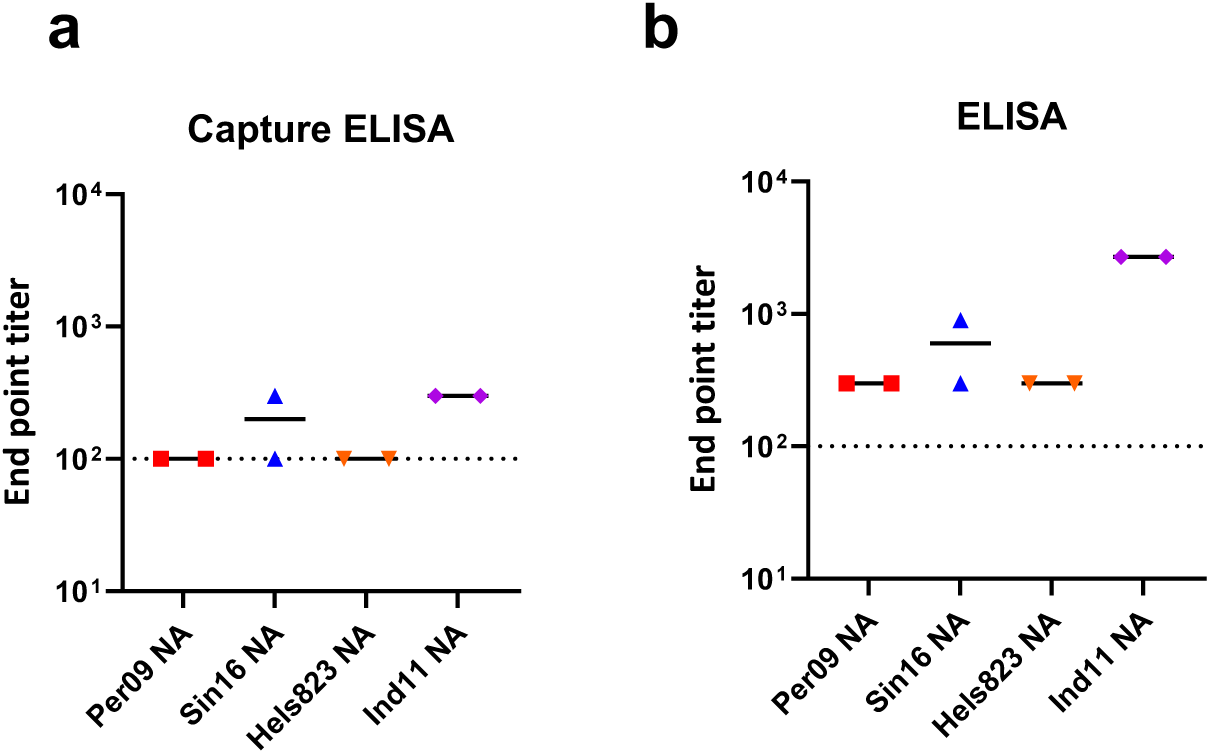
IgG titers against tetrabrachion zipper domain are induced by tetNA immunization. Tetrabrachion domain fused to human serum albumin was captured in nickel coated (**a**) or adsorbed in conventional ELISA plates (**b**). IgG titers in Per09, Sin16, Hels823, and Ind11 NA immune serum were determined using 3-fold serial dilutions and expressed as endpoint titers. Dots represent experimental duplicates of the pooled sera. Dashed lines represent the limit of detection.

**Supplementary figure 3.**
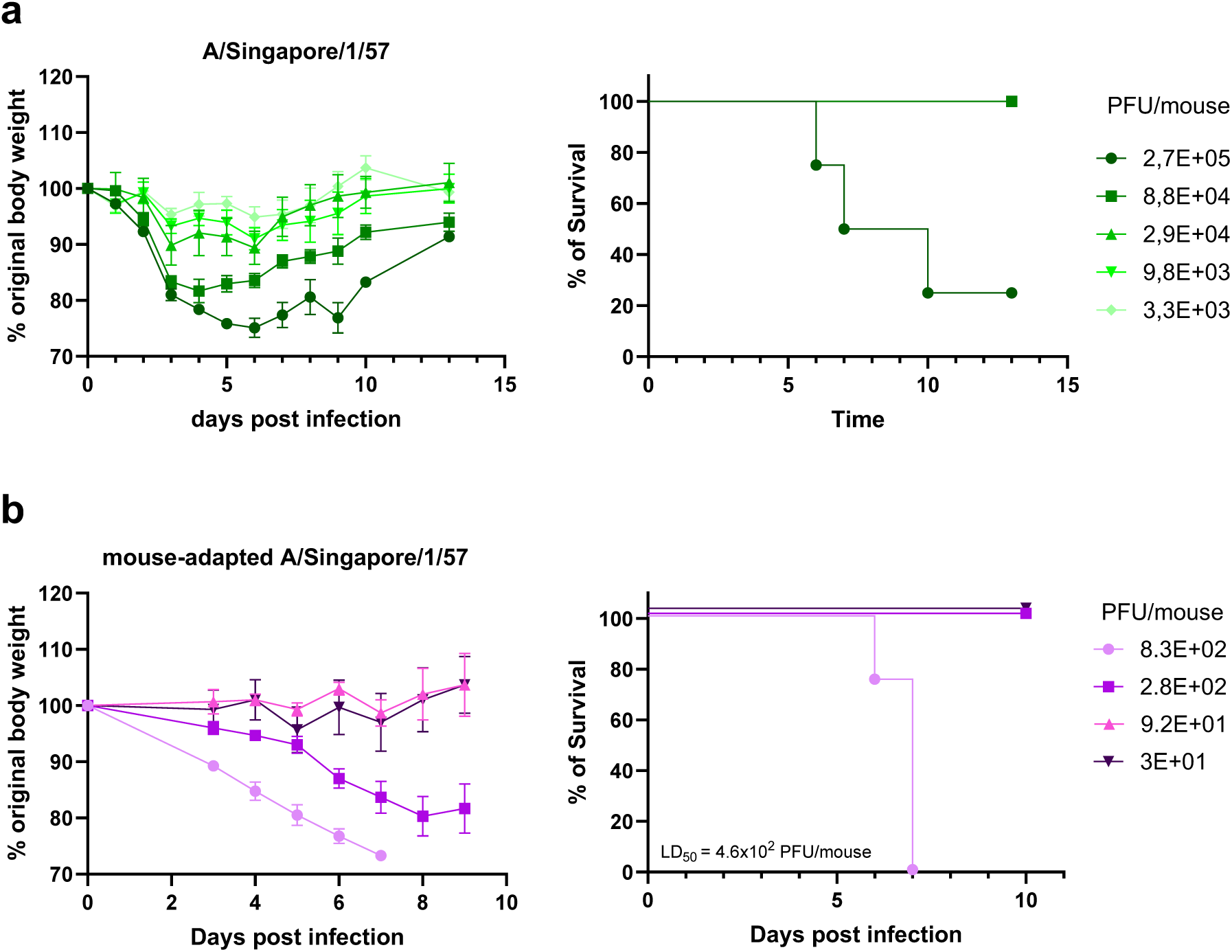
Pathogenicity of A/Singapore/1/1957 in DBA/2J mice. (**a**) Relative weight and survival of mice inoculated with a 3-fold serial dilution of parental Sin57. (**b**) Relative weight and survival of mice inoculated with a 3-fold serial dilution of mouse-adapted A/Singapore/1/1957.

**Supplementary figure 4.**
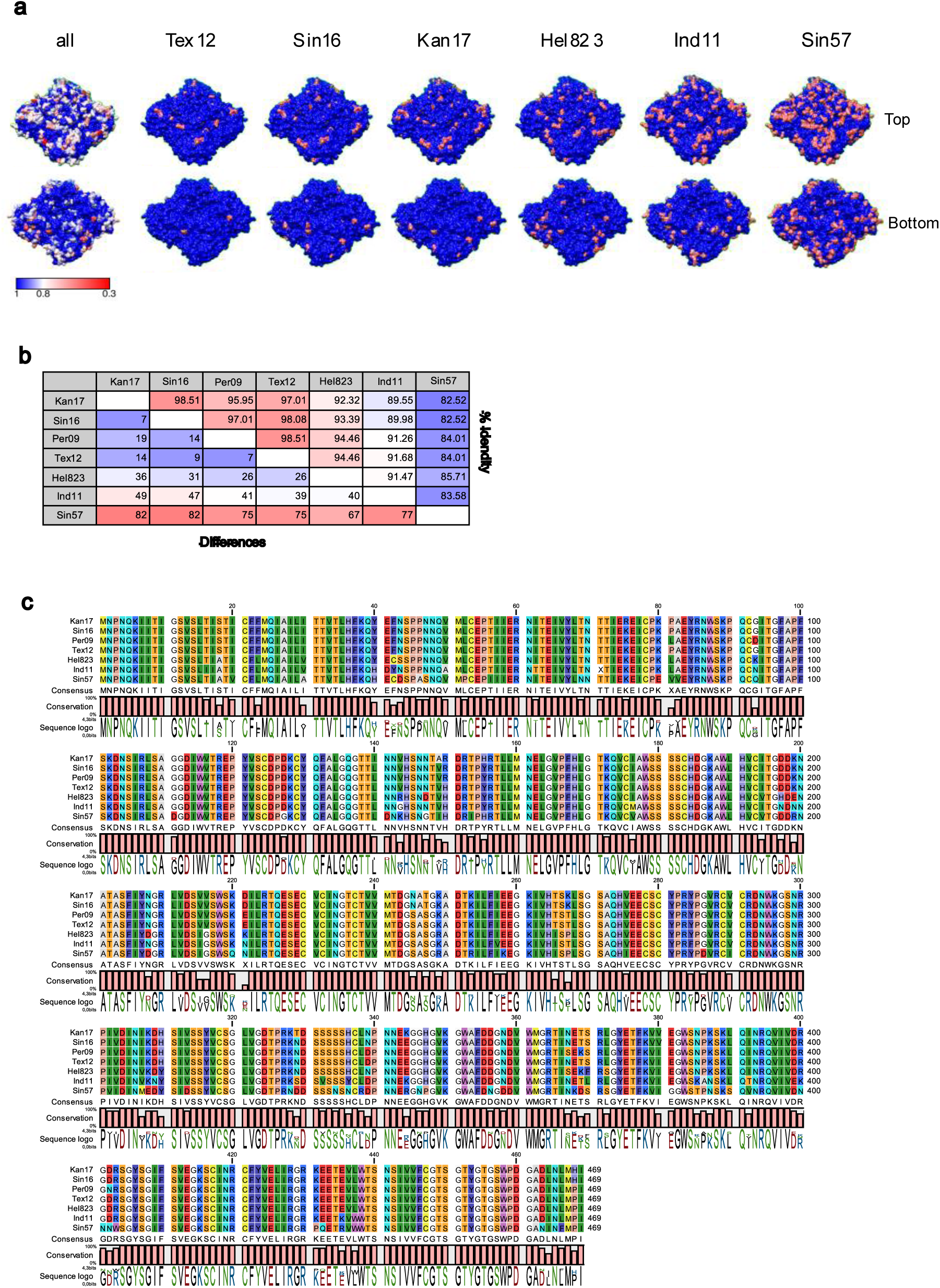
Amino acid conservation among N2s used in the study. (a) Conservancy among All N2s, Tex12, Sin16, Kan17, Hel823, Ind11 and Sin57 relative to NA Per09 shown on surface representation of A/Perth/16/2009 N2 (PBD ID: 6BR5). (b) Pairwise amino acid sequence identity matrix indicating the percentage identity and differences. (c) Amino acid sequence alignment.

